# Characterization and Engineering of *Streptomyces griseofuscus* DSM 40191 as a Potential Host for Heterologous Expression of Biosynthetic Gene Clusters

**DOI:** 10.1101/2020.11.06.372458

**Authors:** Tetiana Gren, Christopher M. Whitford, Omkar S. Mohite, Tue S. Jørgensen, Eftychia E. Kontou, Julie B. Nielsen, Sang Yup Lee, Tilmann Weber

**Affiliations:** The Novo Nordisk Foundation Center for Biosustainability, Technical University of Denmark, Denmark; Metabolic and Biomolecular Engineering National Research Laboratory, Department of Chemical and Biomolecular Engineering, Center for Systems and Synthetic Biotechnology, Institute for the BioCentury, Korea Advanced Institute of Science and Technology, Republic of Korea

## Abstract

*Streptomyces griseofuscus* DSM 40191 is a fast growing *Streptomyces* strain that remains largely underexplored as a heterologous host. Here, we report the genome mining of *S. griseofuscus*, followed by the detailed exploration of its phenotype, including production of native secondary metabolites and ability to utilise carbon, nitrogen, sulphur and phosphorus sources. Furthermore, several routes for genetic engineering of *S. griseofuscus* were explored, including use of GusA-based vectors, CRISPR-Cas9 and CRISPR-cBEST-mediated knockouts. Using CRISPR-BEST technology, core genes of 4 biosynthetic gene clusters (BGCs) that are situated on the chromosome arms were inactivated and the outcomes of the inactivations were tested. Two out of the three native plasmids were cured using CRISPR-Cas9 technology, leading to the generation of strain *S. griseofuscus* DEL1. DEL1 was further modified by full deletion of a pentamycin BGC and an unknown NRPS BGC, leading to the generation of strain DEL2, lacking approx. 500 kbp of the genome, which corresponds to a 5,19% genome reduction. Sequencing confirmed that DEL2 does not bear any crucial off-target effects or rearrangements in its genome. It can be characterized by faster growth and inability to produce three main native metabolites of *S. griseofuscus*: lankacidin, lankamycin, pentamycin and their derivatives. To test the ability of DEL2 to heterologously produce secondary metabolites, the actinorhodin BGC was used. We were able to confirm the production of actinorhodin by both *S. griseofuscus* wild type and DEL2. We believe that this strain will serve as a good chassis for heterologous expression of BGCs.

**Importance:** The rise of antibacterial resistance calls on the development of the next generation of antibiotics, majority of which are derived from natural compounds, produced by actinomycetes. The manipulation, refactoring and expression of BGCs coding for such natural products is a promising approach in secondary metabolite discovery. Thus, the development of a versatile panel of heterologous hosts for the expression of BGCs is essential. We believe that first-to-date systematic, detailed characterisation of *S. griseofuscus*, a highly promising chassis strain, will not only facilitate the further development of this particular strain, but also will set a blueprint for characterisation of other potential hosts.

## Introduction

The vast majority of clinically used antibiotics, antifungals, anticancer and immunosuppressive drugs are derived from natural products (1), originating from soil-inhabiting Actinobacteria (2). Genes, necessary for the production of a particular compound, are typically arranged in biosynthetic gene clusters (BGCs) that can be detected using genome mining tools like antiSMASH (3). With the revolution in whole genome sequencing it became clear that the majority of actinobacterial strains possess genomes of large sizes, with 20 to 40 of BGCs on average (4). However, typically these bacteria only produce few compounds under laboratory conditions (5). Other antibiotics are not produced or produced only in small amounts and require induction by various, frequently unknown, environmental cues (5, 6). In order to unravel the biosynthetic richness of Actinobacteria, multiple genetic engineering methods, e.g. induction of BGCs by co-expression of regulatory genes (7, 8), use of transcription factor decoys (9), refactoring of the BGCs (10, 11) or knockouts of the core genes of “constitutively” produced compounds (12), have been explored. Recently, broader attention has been paid to development of heterologous expression hosts for discovery and characterization of novel metabolites. Heterologous hosts are typically well-characterized strains, that possess plurality of needed characteristics, e.g. fast and disperse growth, amenability to genetic engineering, high yields of produced secondary metabolites (11, 13, 14). The use of such chassis strains is highly advantageous, when the native producer strain can not be genetically engineered or is slow growing (13). At the moment, several of such *Streptomyces* hosts are available, such as derivatives of *S. coelicolor, S. lividans, S. avermitilis, S. chattanoogensis, S. albus, S. venezuelae* (15–20). Recently, several groups have reported the generation of *S. albus* J1074 derivatives that harbor knockouts of multiple native clusters (21, 22) and were successfully used for expression of BGCs (23, 24). The reported success rate of BGCs expression, however, remains low, reaching 30% (14). We believe that the further extension of the panel of heterologous hosts will be highly beneficial to solve this challenge.

One of the promising potential heterologous hosts, *S. griseofuscus*, was first isolated in Japan and used as a host for isolation and propagation of bacteriophages and plasmids (25–28). Some of the strains were used for industrial production of ε-poly-L-lysine (29–31) and puromycin (32), and reported as a source of novel compounds (9, 33). Even though several publications have stipulated multiple positive characteristics *of S. griseofuscus*, e.g. fast growth, ease of transformation and genetic manipulation, it was never methodically studied regarding its qualities as a potential platform for expression of BGCs.

In this paper we describe a comprehensive genotypic and phenotypic characterization of *S. griseofuscus* DSM40191 as a potential heterologous production host. The *De novo* sequenced genome of *S. griseofuscus* was used for mining of natively encoded BGCs. For the first time, we have explored use of various genetic tools for engineering of *S. griseofuscus*, ranging from the use of integrative and replicative vectors, CRISPR-Cas9 and CRISPR-BEST based knockouts to multiplexed-targeting of several BGCs using one CRISPR-BEST construct. All of the generated knockouts were tested regarding their growth and production of metabolites.

## Results

### 1. Genome mining and comparative analysis

*Streptomyces griseofuscus* DSM 40191 (=NRRL B-5429) was received from the German Collection of Microorganisms and Cell Cultures. We re-sequenced and assembled the genome of *S. griseofuscus de novo* (34). It consists of a linear 8,721,740 bp chromosome and three plasmids: pSGRIFU1 (220kb), pSGRIFU2 (88kb), and pSGRIFU3 (86kb) (Fig. 1).

**Fig 1.**
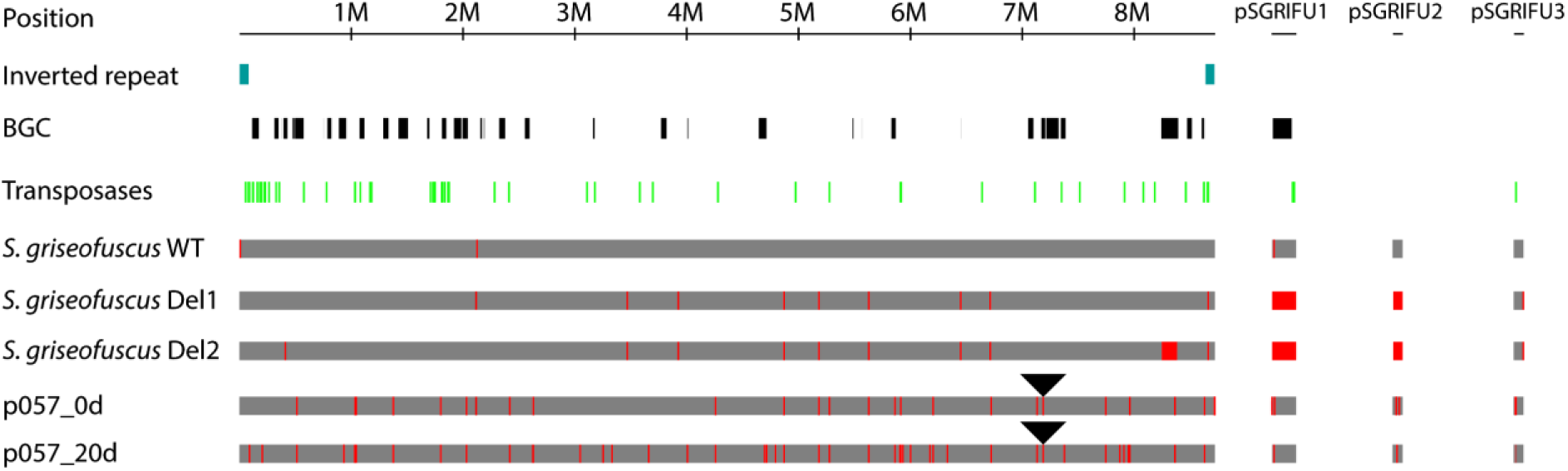
Overview of *S. griseofuscus* strains and mutations. The positions of genomic inverted repeats are highlighted in blue, BGCs in black and transposases in green. The positions of identified mutations are highlighted in red. Mutations detected in the wild type Illumina dataset can be considered technical noise. Position of the CRISPR-cBEST introduced STOP-codon is indicated with a black triangle. Strains p057_0D and p057_20D relate to the long term cultivation experiment, in which CRISPR-cBEST generated strain *S. griseofuscus* IHEP81_06602 (p057_0D), that contains an introduced STOP-codon in BGC 30, was transferred 20 consecutive times in liquid ISP2 media without selective pressure, thus generating strain p057_20D.

#### Whole genome comparison between *S. griseofuscus* DSM40191, *S. coelicolor* A3(2), *S. venezuelae* ATCC 10712

Here, we analyzed how much genomic content is shared between *S. griseofuscus* DSM40191 and other well-studied model *Streptomyces* strains *S. coelicolor* A3(2) and *S. venezuelae* ATCC 10712. Their genomes were downloaded from NCBI (accession IDs NC_003888 and NZ_CP029197) and compared to *S. griseofuscus* DSM 40191 by calculating bidirectional best blastp hits between the genes. We found that 3918 genes were shared across all three genomes (core genome), whereas *S. griseofuscus* shared additional 937 and 522 genes with *S. coelicolor* and *S. venezuelae* respectively (Fig. 2A, Dataset 1). The net total of genes present across the three strains (pangenome set) was 13415. In order to understand the biological functions of the shared genes, we annotated all three genomes with KEGG biological subsystems (35). For the genome of *S. griseofuscus*, we found 2842 ortholog genes in KEGG, which are involved in 1498 KEGG reactions. Whereas the genomes of *S coelicolor* and *S. venezuelae* contained 8152 and 7112 genes, which map to 2900 and 2670 KEGG gene IDs, and 1475 and 1452 KEGG reaction IDs, respectively (Dataset 1). After comparing KEGG genes and reactions content, we found that the three genomes shared 1488 common KEGG gene IDs and 1280 reactions. Next, we compared the distribution of number of genes and number reactions belonging to different KEGG pathways across three organisms (Fig. 2C & D). We found that the number of genes involved in subsystems such as membrane transport, signaling and cellular processes were significantly lower in *S. griseofuscus* than others. Whereas, the number of genes involved in the subsystems like metabolism of terpenoids and polyketides, genetic information processing, amino acid metabolism, xenobiotics biodegradation and metabolism were significantly larger in *S. griseofuscus* (Fig. 2C). Comparison of number of reactions across different metabolic pathways showed that the three strains shared very similar metabolism. However, the number of reactions involved in pathways such as amino acid metabolism was still larger in *S. griseofuscus* (Fig. 2D). Overall, we found that *S. griseofuscus* and *S. coelicolor* had larger genomic and metabolic content than *S. venezuelae*. Additionally, higher genomic and metabolic contents were shared between *S. griseofuscus* and *S. coelicolor*. In the later section, we compared the phenotype microarray data of these strains to get experimental understanding of their metabolic growth capabilities.

**Fig 2.**
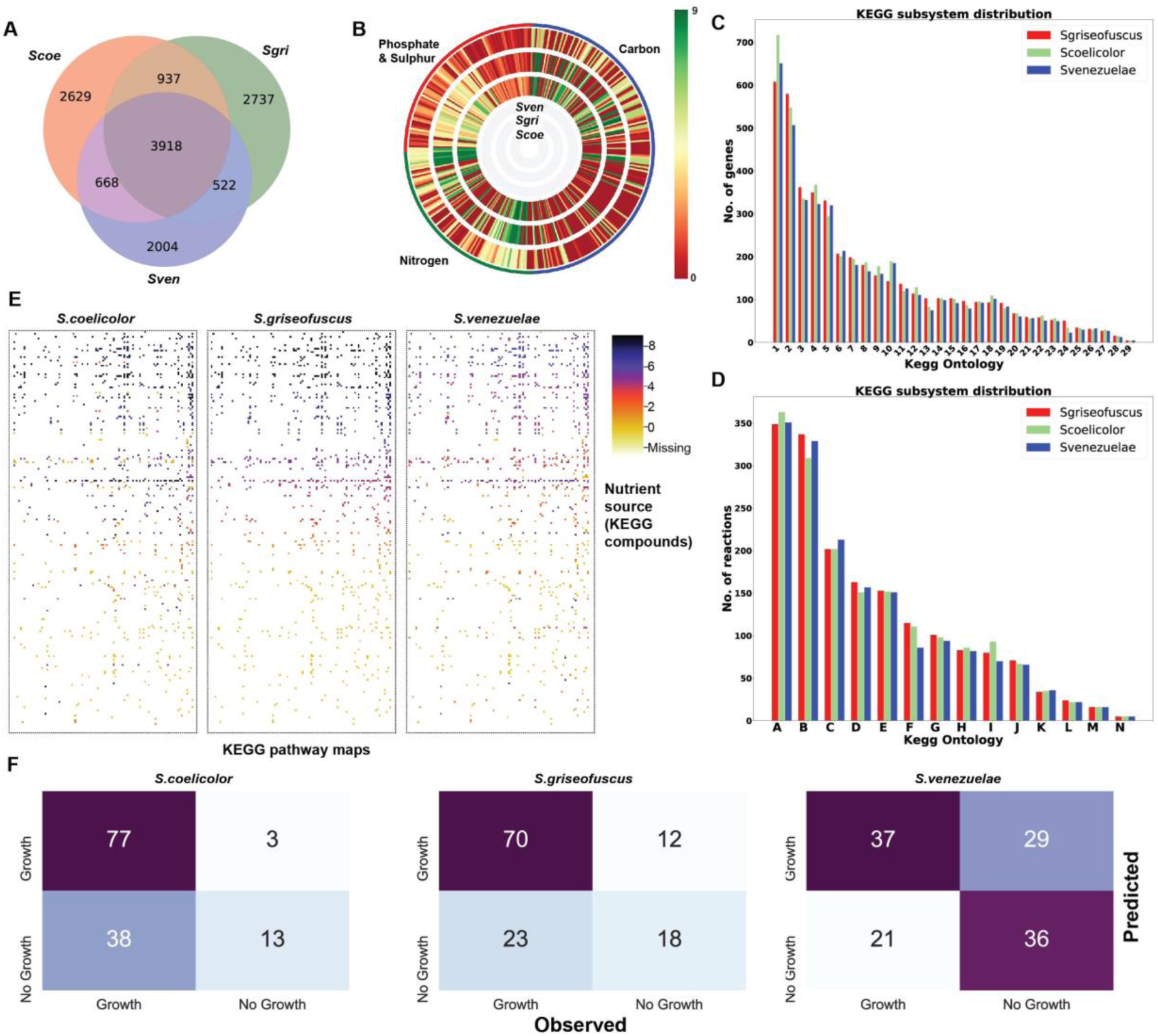
Genome and phenotype microarray comparison of three strains. A. Number of genes shared among the three organisms - *S. griseofuscus* (sgri), *S. coelicolor* (scoe), *S. venezuelae* (sven). B. Phenotype microarray data represented by activity index rings generated using DuctApe across 4 biolog plates consisting 379 carbon, nitrogen, phosphate and sulphate nutrient sources across three organisms. C. Distribution of number of genes per KEGG subsystem across three organisms. For details on this dataset, please see Dataset 1. D. Distribution of number of reactions per KEGG metabolic pathway across three organisms. A-Carbohydrate metabolism; B-Amino acid metabolism; C-Metabolism of cofactors and vitamins, D-Lipid metabolism, E-Nucleotide metabolism, F-Metabolism of terpenoids and polyketides, G - Energy metabolism, H-Biosynthesis of other secondary metabolites, I-Xenobiotics biodegradation and metabolism, J-Metabolism of other amino acids, K-Glycan biosynthesis and metabolism, L-Translation, M-Not included in regular maps, N-Signal transduction.E. Heatmap with activity index of different KEGG nutrients (y-axis) against KEGG pathway maps (x-axis). F. Confusion matrix representing genome-scale model based prediction and observed growth phenotypes of three organisms.

### 2. Phenotype characterization

#### Growth and sporulation on different media

To systematically study a particular strain, it is important to choose an appropriate media that will support fast growth and sporulation in liquid and solid cultures. Therefore, we tested several solid media and have chosen MS as the most accommodating for sporulation (Fig. S1, Table S1). To confirm disperse growth in liquid media, optical density at 600 nm (OD_600_) based growth curve was built in liquid ISP2 (Fig. S2, Table S1).

#### Comparison of physiological features of *S. griseofuscus, S. coelicolor* and *S. venezuelae* using Biolog microarrays

In order to characterize the phenotype of *S. griseofuscus* and its ability to utilize different substrates, we have conducted a multiple parallel cultivation using BioLog microarrays. This technology is not easily applicable for studying actinobacterial strains due to the formation of “clumps’’ of mycelia. However, in the case of *S. griseofuscus*, its uncomplicated growth enables such studies. As a direct comparison, we have used well studied heterologous hosts *S. coelicolor* and *S. venezuelae*. Previously, parallel micro-scale cultivations were used for characterization of an industrially important *S. lividans* TK24 (36).

We tested a total of 379 substrates, including 190 different carbon sources (PM1 and PM2), 95 nitrogen sources (PM3), 94 phosphate and sulphur sources (PM4). The kinetic growth data from Biolog was analyzed together with the genomes using DuctApe software (37) that correlated genomic and phenomic data based on KEGG metabolic pathways. An activity index between 0 to 9 was used to represent the growth on each substrate, where activity index higher than 3 was used as a cutoff to define growth. We found that 171, 172 and 117 of the 379 substrates were utilized by *S. griseofuscus, S. coelicolor* and *S. venezuelae*, respectively (Dataset 2). Comparing the growth of three different strains, we found that 90 substrates were commonly utilized by all three strains, whereas 14, 19 and 7 substrates were utilized uniquely by *S. griseofuscus, S. coelicolor* and *S. venezuelae*, respectively. Some of the substrates uniquely utilized by *S. griseofuscus* include ethanolamine, 2-aminoethanol, cytidine, thymidine, D-serine and D-threonine. Additionally, we found that *S. griseofuscus* shared a total of 145 common growth substrates with *S. coelicolor*, signalling higher mutual metabolic similarity.

Next, we analyzed growth on substrates with different nutrient source categories. We observed that a total of 72 carbon sources were utilized by *S. coelicolor*, which was higher than the number of carbon sources used by *S. griseofuscus* (64) and *S. venezuelae* (61). In particular, *S. coelicolor* could utilize more substrates involved in carbohydrate metabolism. Carbon sources uniquely utilized by *S. griseofuscus* include 2-aminoethanol, alpha-keto-valeric acid, D-malic acid. On the contrary, the number of nitrogen sources utilized were higher in *S. griseofuscus* (60) as compared to *S. coelicolor* (52) and *S. venezuelae* (49). These could be primarily attributed to the categories amino acid metabolism and other non-defined classes of metabolism. Unique nitrogen sources utilized by *S. griseofuscus* include L-phenylalanine, D-serine, ethanolamine. We found that *S. venezuelae* could only use 6 of the phosphate sources which was substantially lower than both *S. griseofuscus* (47) and *S. coelicolor* (46). Uniquely utilized phosphate sources by *S. griseofuscus* included 2-aminoethyl phosphonic acid and dithiophosphate. In general, we observe that the capability of *S. griseofuscus* to utilize different nutrient source categories is much higher than *S. venezuelae*, and is similar or even higher than *S. coelicolor*. Comparison of the ability of *S. griseofuscus* to grow on different nutrient sources can guide the design of growth media and, thus, leads to optimal growth and metabolite production.

To investigate the connection between these growth activity profiles and the genomic diversity of the strains, a matrix was generated using the *dape* module of DuctApe, where the activity on different nutrients (rows) that are part of different KEGG pathways (columns) is highlighted (Fig. 2E, Dataset 3). For example, the average growth activity indices of all the nitrogen source nutrients belonging to KEGG pathway of biosynthesis of amino acids (map:01230) were 6.51, 5.96 and 4.18 for *S. griseofuscus, S. coelicolor* and *S. venezuelae*. In particular, thiamine metabolism pathway (map:00730) showed higher average growth index on nitrogen nutrients in *S. griseofuscus* (7.11) as compared to *S. coelicolor* (5.89) and *S. griseofuscus* (5.78). Overall, the growth activity heatmaps of nutrients vs KEGG pathways were more similar in *S. griseofuscus* and *S. coelicolor*. Whereas, *S. venezuelae* was found to have lower growth activity across nutrients from different pathways. The higher genomic similarity between *S. grisoefuscus* and *S. coelicolor* that was observed in the previous section further corroborates with this phenomic similarity. In addition to this genome to phenome comparison based on KEGG pathways, we used genome-scale metabolic models to compare the *in-silico* predicted growth against observed phenotypes across different substrates (Fig. 2F). We reconstructed draft genome-scale metabolic models for *S. griseofuscus* and *S. venezuelae* based on homology comparison against the genome scale model of *S. coelicolor* (Dataset 4). The models also predicted growth on more number of nutrients in case of *S. griseofuscus* and *S. coelicolor* as compared to *S. venezuelae* (Dataset 5). Thus, we conclude that *S. griseofuscus* possesses very similar or even superior, metabolic capabilities while compared to well studied *Streptomyces* strains.

### 3. Secondary metabolite potential of *Streptomyces griseofuscus*

#### Analysis of the genome using antiSMASH and BiG-SCAPE

In order to estimate the capabilities of the strain to synthesize secondary metabolites, it is important to characterize the BGCs present in the genome. We therefore carried out a genome mining analysis using antiSMASH (3). We detected 35 regions of BGCs encoding for different types of secondary metabolites on the chromosome. These regions can be split into 53 candidate clusters. The megaplasmid pSGRIFU1 (CP051007) includes one antiSMASH-predicted region with seven candidate clusters. No BGCs were detected on pSGRIFU2 and pSGRIFU3. We observed that the genome of *S. griseofuscus* harbored 4 NRPS, 3 PKSI, 5 PKS-NRPS hybrids, 4 other PKS types, 4 terpenes, 4 RiPPs and 11 other types of BGCs as defined by antiSMASH. Some of these BGCs putatively coded for hopene, geosmin, spore pigment, desferrioxamine B, ectoine and pentamycin. Two of the candidate clusters from the plasmid showed similarity to known BGC encoding lankamycin and lankacidin C. We have manually selected some of the candidate clusters for further analysis.

In order to investigate if the BGCs in *S. griseofuscus* are also detected in other *Streptomyces* genomes, we carried out a BGC similarity analysis involving a dataset of 212 publically available complete high-quality *Streptomyces* genomes. In total, 6380 BGCs of different types were detected across this dataset of genomes. We generated a similarity network of 35 regions, 12 manually selected candidate clusters from *S. griseofuscus*, 6380 BGCs from public genomes and 1808 known BGCs from MIBIG database (38) using BiG-SCAPE. The network with the cutoff of 0.3 raw_distance metric was further analyzed using Cytoscape (Fig. 3). All BGC families that did not include one of the BGCs from *S. griseofuscus* were ignored for the subsequent analyses (Dataset 6). We found that only one BGC (region 14) of type NRPS-like was a singleton in the network, uniquely observed in *S. griseofuscus*. We observed that 8 of the BGCs were exclusively present in one other genome, namely *S. rochei* 7434AN4. In addition, further 9 BGCs are also present in *Streptomyces. sp*. endophyte_N2 (Genbank Accn.: CP028719) in addition to *S. rochei* 7434AN4 (Genbank Accn.: AP018517). This suggests that these 17 BGCs from *S. grieseofuscus* are also very rarely observed across streptomycetes. Among BGCs that are relatively common across the dataset, we found that candidate cluster 50 of region 33 had similar BGCs across 18 other *Streptomycetes*, including *S. collinus Tu 365* (Fig. S3), whereas the candidate cluster 51 was similar to known cluster encoding for pentamycin. Overall, we have established that most of the BGCs (33) were also present in *S. rochei* 7434AN4, indicating two genomes with highly similar content.

**Fig 3.**
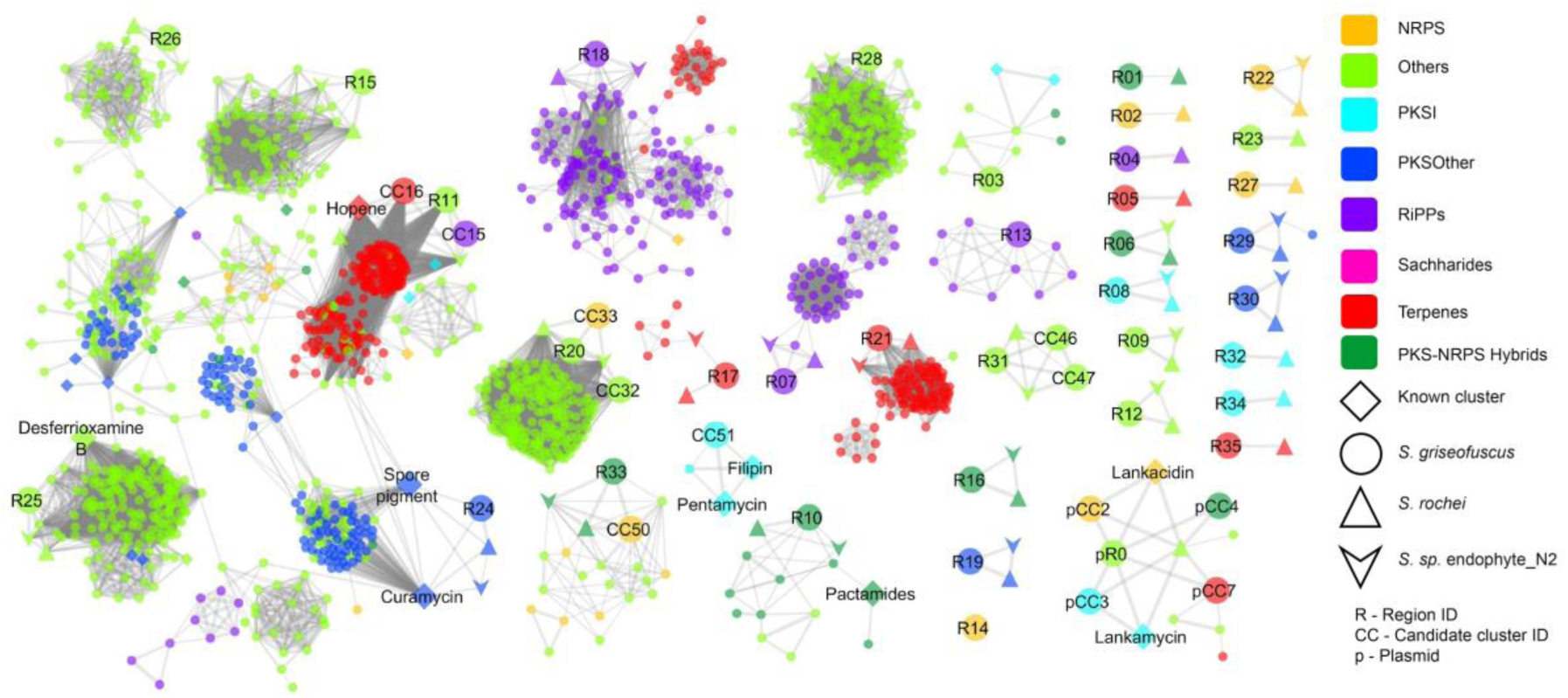
Similarity network of BGCs in *S*.*griseofuscus* against 212 public genomes and MIBIG database of known BGCs. Comparison of all the regions and few selected candidate clusters against BGCs of known compounds from MIBIG database and 212 public genomes. Different colors denote different types of BGCs as shown in legends. BGCs from *S. griseofuscus, S. rochei* 7434AN4, *Streptomyces. sp. endophyte_N2* and MIBIG database are shown with different shapes and sizes. All regions, selected candidate clusters and known BGCs are annotated by text. Detailed comparison of selected BGCs using CORASON can be found in Fig. S3.

This similarity has led us to examine the relation of *S. griseofuscus* to other strains of its species and to *S. rochei* strains. Currently, there are 4 complete assemblies *S. griseofuscus* and 3 of *S. rochei* genomes available in NCBI database. Among these are aforementioned *S. rochei* 7434AN4 and type strain *S. rochei* NRRL B-2410. By calculating pairwise Average Nucleotide Identity (ANI) between all genomes (39), we have identified ANI between *S. griseofuscus* DSM40191 and *S. rochei* 7434AN4 to be at 99.54%, while similarity of *S. rochei* 7434AN4 to the type strain *S. rochei* NRRL B-2410 is at 84.03%, highly similar to the one between *S. rochei* NRRL B-2410 and *S. griseofuscus* DSM 40191 of 83.95%. This clearly signals that *S. rochei* 7434AN4 strain was probably misclassified and is indeed a *S. griseofuscus* strain. This may explain the large number of similar BGCs, shared between *S. griseofuscus* DSM 40191 and *S. rochei* 7434AN4 and high level of similarity between two of the largest plasmids in both strains pSGRIFU1 and pSLA2-L.

#### Characterization of secondary metabolites, produced by the *S. griseofuscus*

In the genome mining study, we have identified several known BGC, studied in other strains. Among these are lankacidin and lankamycin BGCs, encoded on plasmid pSGRIFU1, previously studied in *S. rochei* (40, 41) and pentamycin BGC33 (42). Due to the good preservation of these BGCs in the genome we expected to detect the production of lankamycin, lankacidin-family compounds and pentamycin. In addition, it was reported previously, that some strains of *S. griseofuscus* are able to produce azinomycins A and B (43), acetylcholine esterase inhibitor physostigmine (44), ϵ-poly-L-lysine (29, 30), lankacidin C and A (25). In order to check the production in *S. griseofuscus* DSM 40191, we have performed exploratory cultivation in 5 different liquid media (ISP2, MAM and CDMZ medium (44), minimal medium (MM) (45), and medium 65) (46), that were described in literature for the production of respective compounds and have attempted to identify them in extracts. Lankacidin A, C, and lankamycin were tentatively identified by HR-MS. The production of pentamycin was identified by HR-MS and confirmed using the pentamycin standard (Fig. S4). Physostigmine was not detected in any of the conditions (Table S2).

We have noted the production of hydrophobic extracellular vesicles by *S. griseofuscus*, a widely spread, but poorly studied phenomenon among Actinobacteria (47). It is known that the extracellular vesicles might contain secondary metabolites (48, 49). To study the profile of the extracellular vesicles in *S. griseofuscus*, they were collected and directly injected for LC-MS measurements. Among many compounds we have tentatively identified lankamycin, that was previously detected in the cultivation extractions and is likely released as a part of extracellular vesicles.

### 4. Development of genetic engineering methods

Even though the transformation, conjugation, protoplast generation for *S. griseofuscus* were established before, including attempts of genetic engineering (32, 33, 50) it was never systematically tested with different vectors and engineering methods. While generating a heterologous host strain, it is important to have access to the fastest knockout-leading techniques that lead to the least off-target modifications.

#### Transfer of integrative and replicative GusA-based vectors

As the first step we tested whether *S. griseofuscus* is compatible with *gusA* reporter system plasmids (51): pSETGUS, an integrative phiC31-based plasmid, and pKG1139, a replicative plasmid. Both plasmids were successfully conjugated into *S. griseofuscus* and allowed for visual screening of the exconjugant colonies (Fig. S5). To determine the position of the pSETGUS integration site, which is of importance to rationally utilize it for the integration of desirable elements, we randomly picked three independent *S. griseofuscus* pSETGUS colonies and sequenced them using Oxford Nanopore sequencing, similarly to (52). The exact location of the integration site is at 4,242,328 bp in the HEP81_03793 gene, coding for putative chromosome condensation protein. The determined *attB* site of *S. griseofuscus* contains the conserved core “TT” sequence (53).

#### CRISPR-Cas9 mediated gene knockout

CRISPR-Cas9-based molecular tools offer precision and ease in handling in comparison to other techniques. Over the recent years CRISPR tools have been adapted for use in streptomycetes (54). As the introduction of double strand breaks can lead to rearrangements and off-target effects in the genome, we validated various CRISPR-Cas9-based engineering methods by targeting genes on the chromosome and on one of the plasmids. For this purpose we used pGM1190-based CRISPR-Cas9 plasmid (55), based on a temperature sensitive replicon, shown to be functional in *S. griseofuscus* by using GusA-based pKG1139.

As the first target, we wanted to eliminate plasmid pSGRIFU1 that harbours 4 BGCs, among them lankacidin, lankamycin, cryptic polyketide and carotenoid BGCs. This plasmid has a very high similarity to the plasmid pSLA2-L of *S. rochei*, where these clusters were characterized (41). A sgRNA was designed to target the DNA primase/helicase-coding region, which is essential for plasmid replication. Three random colonies were selected after the CRISPR procedure and sequenced via Illumina whole genome sequencing. Surprisingly, in all clones, both the targeted pSGRIFU1, but also pSGRIFU2, were lost, leaving only plasmid pSGRIFU3 present in the genome. To estimate the amount of changes in the plasmid-cured strains in comparison to the wild type genome, the WGS data was analyzed with breseq, which identified 11 mutations (six SNVs, three insertions, and two deletions). One of the colonies was selected for further work and named DEL1.

In parallel, we attempted to knockout chromosomally located BGC region number 33, which encodes a putative pentamycin BGC and an uncharacterized NRPS BGC (Fig. S3). The conjugation of the knockout plasmid resulted in less than 10 colonies, 2 of which were selected for Illumina MiSEQ sequencing. It revealed that even though both clones accumulated several SNPs, they did not contain the intended mutation (data not shown). Even after the experiment was repeated, we were not able to select knockout-carrying colonies.

In order to verify whether the deletion of pentamycin-NRPS clusters is possible in the plasmid-cured conditions, knockout plasmid was transferred to DEL1. In contrast to the experiments with the wild type, a large number of exconjugants was received. After the plasmid curing, three of the independently received colonies were sequenced with Illumina NextSeq and one of them was additionally sequenced using Nanopore technology. This clone, further referred to as DEL2, was confirmed to contain full deletion of pentamycin-NRPS cluster region and contained a comparatively small amount of SNPs (Fig. 1).

In the strain *S. rochei* 7434AN4, which is closely related to *S. griseofuscus*, curing of all three plasmids has been reported to change the topology of the chromosome from linear to circular (41, 56). It is believed that the *tap*-*tpg* gene pair, which encodes for telomere-associated protein and a terminal protein for end patching, located on both pSLA2-L and pSLA2-M plasmids, is responsible for maintaining the linear architecture of chromosome. Because both the genomes and the associated plasmids in *S. rochei* and *S. griseofuscus* are similar, we investigated if the chromosome of *S. griseofuscus* had circularized during the plasmid curing. We therefore sequenced the strain DEL2 with the Nanopore technology. The assembly graph clearly showed a chromosome with inverted repeat consistent with a linear chromosome. In order to verify presence of the *tap-tpg* homologues in the genome of *S. griseofuscus*, a BLAST search was performed against each gene pair from pSLA2-L and pSLA2-M. The homologues of *tapR1*-*tpgR1* and *tapRM*-*tpgRM* were found on all three plasmids of *S. griseofuscus* (Table S3). This fact could explain the preserved linear topology of the DEL2 chromosome. The removal of putative pSGRIFU1 and 2 *tap*/*tpg* homologues, does not lead to the circularisation of the chromosome, because of the remaining homologous genes, present on pSGRIFU3.

Both of the DEL1 and DEL2 strains did not show any significant changes in their morphology, growth or sporulation (Fig. S1, S2, Table S1). In order to verify the influence of genetic manipulations on the metabolites produced by DEL1 and DEL2, parallel cultivations in ISP2 media were made. In comparison to the wild type, strain DEL2 lost the possibility to produce pentamycin, lankacidins and lankamycin, as it was expected (Fig. S4).

In order to test whether *S. griseofuscus* is suitable for the expression of heterologous BGCs, we have expressed *S. coelicolor* actinorhodin BGC in the wild type and DEL2. As evident from the formation of dark-blue halo, wild type and DEL2 strains are both able to produce actinorhodin in heterologous conditions (Fig. S6).

#### CRISPR-cBEST mediated knockouts

The recent extension of our CRISPR-Cas9 toolkit, CRISPR-cBEST system (57), utilizes cytidine deaminase, fused to dCas9. It allows for introduction of STOP-codons in the positions of interest by converting CG base pairs to AT. Recently, we have reported the use of this system in *S. griseofuscus* (57). In order to test the usability of CRISPR-cBEST for engineering of *S. griseofuscus*, the targeted BGCs were selected on the so-called “arm” regions of the chromosome (57). It is well known that the introduction of double strand breaks of the DNA by Cas9 might lead to multiple unwanted consequences. It is particularly dangerous in case of the ends of the chromosome, that by itself frequently undergo rearrangements (58). Therefore, these BGC regions are particularly difficult to engineer. In order to verify whether CRISPR-cBEST system would help to omit these limitations, the targets were selected in 4 different BGCs-containing regions number 4, 30, 31, 34. on the right and left arms of the chromosome. The pCRISPR-CBE plasmids were constructed according to the protocol (59) sequenced and transferred to *S. griseofuscus* via conjugation. Correct clones with the STOP-codons in BGC regions 4, 30, 31 and 34 were confirmed via Sanger sequencing of the region of interest (57). In order to determine the outcomes of each mutation, the morphology, growth and metabolite production was assessed and individually described (Fig. S1, S2 Table S1). We have grown all of the CRISPR-cBEST generated mutants in ISP2 liquid media and compared their production profiles to the wild type (data is not shown). In these initial tests, we were not able to identify specific metabolites, produced from each of these BGCs, possibly because production conditions for these metabolites were not met, or they are cryptic.

It was shown that by using the multiplexed CRISPR-cBEST plasmids it is possible to target multiple genes from different BGCs in *S. coelicolor* (57). Therefore, it was our next target to verify such a possibility in *S. griseofuscus*. For this purpose, a multiplex plasmid was constructed, targeting 4 BGCs on the left arm of the chromosome. The sgRNA guides, selected earlier, were used, yielding plasmid pCRISPR-MCBE-1-2-4-6, targeting BGCs 1, 2, 4 and 6. The plasmid was verified via Sanger sequencing and transferred to *S. griseofuscus* via conjugation. Up to 24 exconjugant colonies were tested via PCR. Each of the targeted regions was amplified using a selected set of primers, the fragments were purified and sequenced by Sanger sequencing. As the result of the screening, for each of the targeted regions at least one successful editing event was detected. We were able to select a colony of *S. griseofuscus* pCRISPR-MCBE-1-2-4-6 with a total of three edited targets (E3I2) (Table S4). Strain E3I2 has exhibited signs of sporulation deficiencies and changes in morphology, that might be related to the specific combination of the mutations that were introduced (Fig. S1). However, growth of this strain was clearly not inhibited in liquid cultures (Fig. S2, Table S1). In addition, the metabolite biosynthesis profile of E3I2 was verified in ISP2 liquid media (data is not shown). We were not able to identify specific metabolites, linked to the inactivated BGC, probably because the conditions for the production of these metabolites were not met or these particular BGCs were not expressed.

One of the significant problems for CRISPR-Cas9-mediated targeting are unwanted off-target effects. Similarly, such problems exist while using CRISPR-BEST systems. It was shown that while using the CRISPR-BEST for the generation of knockouts in *S. coelicolor*, a relatively small amount of SNPs can be observed (57). However, the influence of the presence of CRISPR-BEST plasmid on the accumulation of the SNPs over the time during continuous cultivation was never studied.

In order to study these effects we have performed a long term cultivation experiment with CRISPR-cBEST generated mutant strain *S. griseofuscus* HEP81_06602 (p057). The initial and resulting strains were sequenced using Illumina NextSEQ and compared to the wild type strain, using breseq analysis (Fig. 1 and 4). Notably, the introduced Trp221Stop mutation in the putative gene HEP81_06602 (BGC 30) was maintained even after 20 transfers without the antibiotic pressure (Fig. 1).

**Fig 4.**
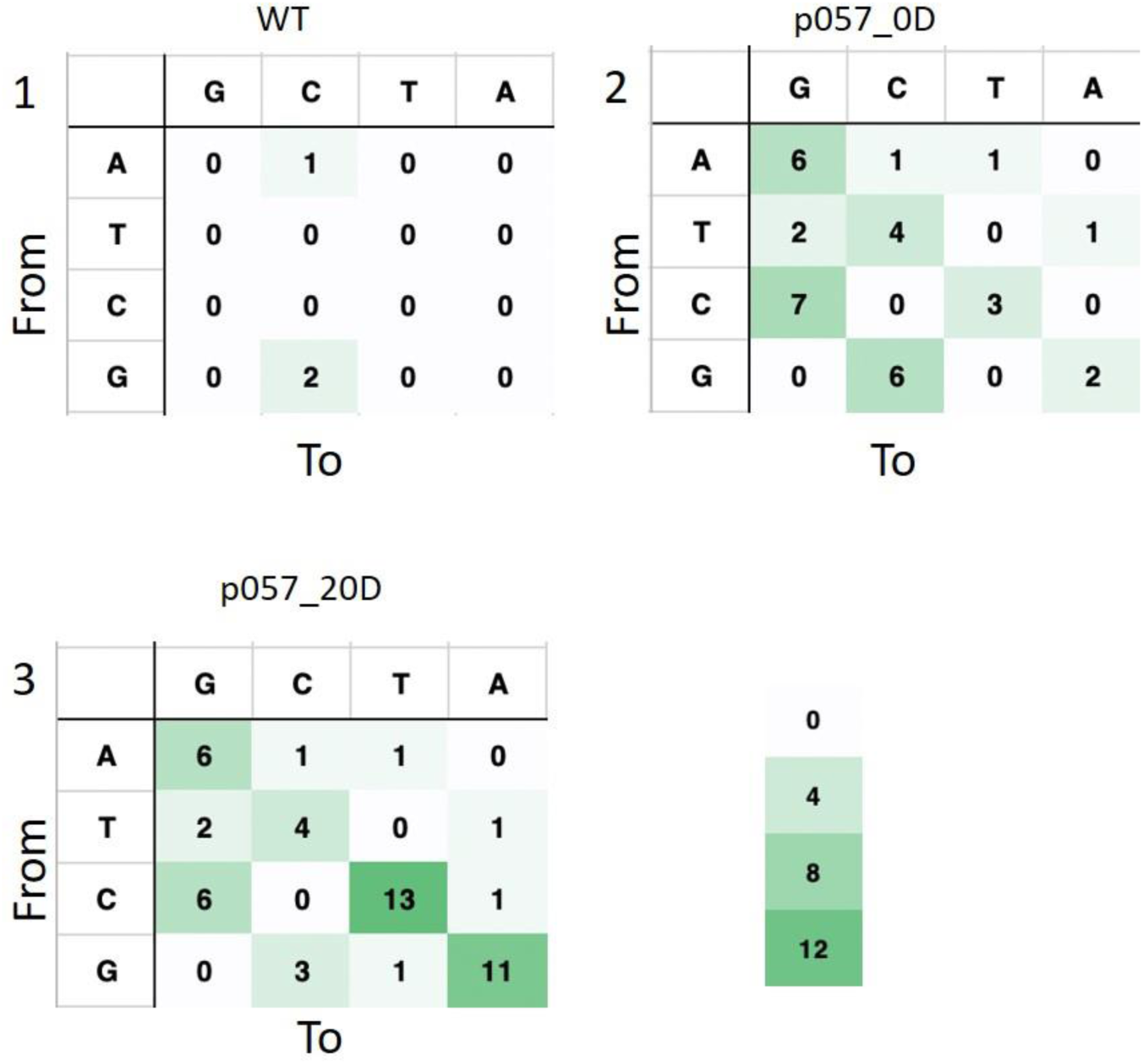
Genome wide off-target evaluation of CRISPR-BEST mediated mutations in the strain preserved and sequenced after introduction of Stop-codon (p057_0D; 2) in comparison to the same strain that was passed consecutively 20 times in liquid cultures (p057_20D; 3). Mutations, predicted for the WT Illumina dataset (1), used to produce the reference, the level of which thus can be considered a technical noise.

The breseq analysis of the Illumina-generated reads of the wild type genome had detected 3 SNPs, which can be considered a technical baseline. In case of strain p057_0D this number increased to 33 SNPs altogether, with a majority of them being C to T exchanges, which putatively can be attributed to nonspecific activity of the CRISPR-BEST cytidine aminase. After 20 consecutive transfers this number increased to 50 SNPs with a majority of them being C to T and A to G exchanges. Both numbers are falling in the range of the previously reported for *S. coelicolor* (57) and are promising for the engineering of *S. griseofuscus*.

## Discussion

Actinobacterial genomes usually code for 20-30 secondary metabolites (4). However, under laboratory conditions a limited number of compounds are produced, possibly due to the absence of appropriate cues for the BGC expression. One strategy to address this challenge is the expression of BGCs in heterologous hosts. Even though there are several of such strains available, the current rate of successful expression of BGCs is low (14). Here, we present a further diversification of the heterologous strain panel with a potentially highly adaptable, flexible and easy-to-handle heterologous host, *S. griseofuscus*. The literature-provided information regarding *S. griseofuscus* is comparably dated. Therefore, we have decided to start our work with the wild type strain *S. griseofuscus* DSM 40191 and test its qualities first hand. While exploring its phylogenetic position, we have identified an ANI to *S. rochei* 7434AN4 of 99.54%, while the ANI of *S. rochei* 7434AN4 to the type strain *S. rochei* NRRL B-2410 is only at 84.03%. This indicates that *S. rochei* 7434AN4 may be misclassified and rather should be included in the *griseofuscus* group. *S. rochei* 7434AN4 is described in detail as a producer of lankamycins, lankacidins and pentamycin (40, 42). However, it was never systematically characterised regarding its genotype, phenotype or possibility of use of different genetic engineering methods. Therefore, detecting the possible similarity between *S. rochei* 7434AN4 and *S. griseofuscus* DSM40191 has been helpful in order to ease characterisation of *S. griseofuscus* metabolic profile.

The comparison of *S. griseofuscus* genome to genomes of the model streptomycetes *S. coelicolor* and *S. venezuelae* has revealed a close genetic similarity, retained in the ability to process sources of carbon, nitrogen and sulphur. Notably, *S. griseofuscus* is genetically, and phenotypically closer related to *S. coelicolor* than to *S. venezuelae*. This enables the use of the methods developed for *S. coelicolor. S. griseofuscus* utilizes various media sources for its growth, displaying apparent metabolic flexibility, needed for heterologous hosts. In the growth tests, the strain had displayed fast uncomplicated growth in the liquid media, compatible with OD measurements, and abundant sporulation on solid media. In addition, we demonstrated that *S. griseofuscus* produces native metabolites lankacidin, lankamycin and pentamycin, as it was expected from its phylogenetic “proximity” to *S. rochei* 7434AN4. This information was later used in the selection of target BGCs for genetic engineering.

Heterologous host strains have to be frequently manipulated genetically in order to be molded as producers of specific secondary metabolites. Therefore, it is important to have access to a set of efficient genetic engineering techniques. We have tested various methods, such as integrative and replicative vectors, CRISPR-Cas9 based knockouts and CRISPR-cBEST base editing. *S. griseofuscus* is accessible to all these methods, implemented for single or multiplexed knockouts, which generate minimal off-target effects. Even though CRISPR-based systems were shown to be applicable in Actinobacteria, their use is still largely limited to model strains and typically requires further adjustments (60). It is, therefore, highly beneficial that *S. griseofuscus* does not require any of such additional efforts.

Our first target for the genome reduction of *S. griseofuscus* has been the curing of the largest plasmid pSGRIFU1, highly similar to pSLA2-L of *S. rochei* (42). It carries 4 BGCs, among these the lankacidin and lankamycin BGCs, products which were detected in the wild type supernatant. As it was expected (40), CRISPR-Cas9 mediated curation of pSGRIFU1 led to the disappearance of these metabolites in the supernatants of the mutant strain DEL1. To generate strain DEL1 we have targeted the DNA primase/helicase-coding region of the pSGRIFU1 plasmid, which also has caused the curing of plasmid pSGRIFU2. This could be explained by the presence of several putative off target sites in pSGRIFU2. Curation of pSGRIFU2 was unexpected, but a positive outcome for the genome minimization and did not lead to chromosome cyclisation of DEL1, as it was shown for *S. rochei* (56). We believe that the homologues of genes *tap*/*tpg*, that were detected on plasmid pSGRIFU3, are needed to sustain the architecture of the chromosome.

Furthermore, we have successfully removed pentamycin and NRPS BGC from the genome of DEL1, generating strain DEL2. The metabolomic studies have shown that strain DEL2 does not produce pentamycin (Fig. S5).

DEL2 can be considered as the first genome reduced strain of *S. griseofuscus* with 5,19% of its genomes removed without any negative effects on its growth, morphology or sporulation levels. In addition, actinorhodin BGC was successfully expressed in DEL2 (Fig. S6). Therefore, we believe that strain DEL2 can be considered as the first step towards developing heterologous production host, suitable for expression of polyketide BGCs.

## Conclusions

For the last several decades, *S. griseofuscus* has remained largely unexplored. In this paper, a detailed characterization of *S. griseofuscus* DSM 40191 is presented, including studies of its genotype, phenotype and metabolic capabilities. We believe that the results, described in this manuscript clearly demonstrate the high potential of *S. griseofuscus* strain as to its further development as heterologous host strain. We plan to continue our work on the genome reduction of *S. griseofuscus* DEL2, using step-by-step removal of BGCs with simultaneous characterisation of mutants growth and metabolite production. In the following next phase of development, we plan to adapt the resulting strain to expression of different types of BGCs, by introducing additional integration sites and by modifying the primary metabolism to suit the precursor needs.

## Materials and Methods

### 1. Strains, used or constructed in this study

Strains *S. griseofuscus* DSM 40191 and *S. venezuelae* DSM 40230 were received from DSMZ strain collection as freeze-dried pellets. Strain *S. coelicolor* M145 was received from Y. Tong (NBC group, DTU, Denmark). *E. coli* Top10 and Mach1 (both from Thermo Fisher Scientific) were used as general cloning hosts in this study. Other strains that were used in this study are described in the Table S4.

### 2. Growth and cultivation conditions

All of the Streptomyces strains were cultivated using 3 different solid media: MS (45), ISP2 and ISP4 (International Streptomyces Project medium 2 and 4; premixed Difco™ ISP2 dehydrated medium (Fisher Scientific, #DF0770-17-9) and premixed Difco™ ISP4 dehydrated medium (Fisher Scientific, #DF0772-17-7)). DSMZ medium 65 (GYM *Streptomyces* medium) is the medium recommended by the German Collection of Microorganisms and Cell Cultures (DSMZ) for cultivation of *S. griseofuscus* DSM 40191 and prepared as described by DSMZ (46). CDMZ, a chemically defined media for detection of physostigmine, was prepared as described in (44). Otherwise, *Streptomyces* minimal media (45), MAM, molicidin A medium, and version of ISP2 without addition of agar, were used for liquid media cultivations. For cultivations in liquid medium 250 ml shake flasks were used in combination with approximately 30 3 mm glass beads, incubated at 30 °C and 200 rpm. For the cultivations in liquid media, a two stage cultivation principle was used. At first, pre-cultures were directly inoculated with spore suspensions, that were collected from fully grown MS plates and incubated overnight. The OD600 was measured for each pre-culture and re-inoculated in the main culture up to the final OD600 of 0.1. The OD600 measurements were carried out in the 1.5 mL disposable polystyrene spectrophotometer cuvettes. Before each measurement, the cuvettes were thoroughly mixed using a vortex-mixer and each cuvette was measured three times. Therefore, each presented data point corresponds to a total of nine measurements. The maximum specific growth rate µ_max_ was determined by plotting the natural logarithm of all OD_600_ values against the cultivation time. Further on, a linear fit for data points corresponding to the exponential phase was calculated and the slope of that fit corresponds to µ_max_. Using µ_max_, the minimal time needed for the OD_600_ value to double (td) was calculated as described in (61).

### 3. Cloning, primers and constructed plasmids

Spacer sequences, primers, constructed and used plasmids are listed in Table S5. All spacer sequences were selected with the help of CRISPy-web (62, 63).

The procedure of ssDNA oligo bridging was used for integration of spacers into CRISPR plasmids. The CRISPR plasmid of interest was digested with NcoI at 37 °C for 30 min and dephosphorylated using FastAP at 37 °C for another 30 min. The reaction was inactivated by incubation at 65 °C for 10 min. The 20 nt spacers were ordered as oligos from IDT with 20 nt overlaps to the backbone on both sides (Table S5). After dilution to 100 µM, the oligos were diluted to a concentration of 0.2 µM using 1x NEBuffer 2. A 10 µl reaction mix was prepared containing 30 ng of the linearized backbone, 5 µl of the 0.2 µM oligo, and ddH_2_O to 10 µl. 10 µl of NEBuilder^®^ HiFi DNA Assembly Master Mix were added and the reaction was incubated for 1 h at 50 °C. Up to 5 µl were transformed into electrocompetent *E. coli* Mach1. Positive colonies were identified by colony PCR and running the samples on a 2 % agarose gel. Putative positive colonies were further analyzed by the in-house Sanger sequencing. The construction of CRISPR-Cas9 plasmids and their transfer was carried out as described in (59), All *E*.*coli-Streptomyces* conjugation experiments were conducted according to the modified protocol from (45) in which the heat shock step was completely omitted. The MS media with addition of magnesium chloride solution (45) was used for plating of conjugation mixes.

### 4. Genome comparison of *S. griseofuscus, S. coelicolor and S. venezuelae*

To compare genetic features of *S. griseofuscus* with other well studied *Streptomyces* strains, we downloaded the genomes of *S. coelicolor* A(3)2 (NCBI accession.: NC_003888) and *S. venezuelae* ATCC 10712 (NCBI accession: NZ_CP029197). All three genomes were annotated using the KEGG annotation server that describes the biological subsystem of each gene. The number of genes in different biological subsystems were counted for each of the three genomes. Similarly, to compare the metabolic properties, the number of unique KEGG reactions were counted for different metabolic pathways across the three genomes. The genomic and phenotypic microarray data was analyzed together using the *dape* module of the DuctApe software that is used to correlate genomic data with phenomic data for multiple strains. As part of this software, bidirectional best blast hits (E-value threshold – 1e-10) were calculated to generate the pangenome of the three strains. Based on this pangenome, the number of shared genes, unique KEGG gene IDs, and unique KEGG reaction IDs across the three genomes were calculated.

### 5. BioLog phenotypic microarrays and data analysis

Phenotype MicroArrays were ordered from BioLog, which in addition had provided customized protocol for the cell inoculation and measurements. In this study testing plates PM 1 to 4 were used, out of which PM1 and 2 contain Carbon sources, PM2 - Nitrogen sources, PM4 - Phosphorus and Sulphur sources. All the measurements for all strains were performed in three technical and two biological replicates. *S. coelicolor* M145, *S. venezuelae* DSM 40230 and *S. griseofuscus* DSM 40191 were all grown on MS plates for 6 days until clear sporulation signs appeared. The spores were collected in sterile water and diluted to 80% of cell density using BioLog-supplied turbidimeter (catalogue number: 3531). BioLog redox dye mix G was used for all measurements. The recipe of minimal media and other supplementing resources were received from BioLog. The cells were mixed with the prepared media and inoculated in the PM plates using multichannel pipettes and immediately loaded into the OmniLog instrument for measurements (catalogue number: 91171).

The kinetic data for all testing plates PM1 to PM4 generated by BioLog was further analyzed using DuctApe software (37) along with the genomic data. Activity index between 0 to 9 was assigned for each nutrient source to represent the growth activity on each nutrient source. Average activity index was further calculated for the two replicas that were used to generate phenotype data. The average activity index across 379 nutrient sources across 4 PM plates was visualized using activity index rings for the three different strains (Dataset 2). Sources with differential growth activities were further analyzed. To compare the growth activity with genomic features, a matrix with activity on different KEGG nutrients (rows) against KEGG pathways with the nutrient (columns) was calculated. These matrices were compared using heatmaps where rows and columns are ordered as per *S. griseofuscus* growth activity index and number of reactions per KEGG pathway respectively (Dataset 3).

#### Genome scale model reconstruction

We used Python-based genome-scale model reconstruction tool ‘Genome-scale Modelling with Secondary Metabolism’ (GMSM) that implements bidirectional blastp hits-based homology modelling(unpublished) to automatically reconstruct models for both *S. venezuelae* and *S. griseofuscus*. We used a model of *S. coelicolor* A3(2), iMK120848, as a template for the homology modelling. The in-silico growth predictions were made using flux balance analysis on different simulated mediums using COBRApy. Confusion matrix was created from predicted and observed growth phenotypes to compare the model prediction against Biolog data.

### 6. DNA isolation, sequencing and assembly

As described in (34), DNA from *S. griseofuscus* was purified using the Qiagen Genomic Tip 100 (Venlo, Netherlands) kit. Illumina (San Diego, California, USA) WGS libraries were constructed using the KAPA (St. Louis, Missouri, USA) HYPRplus kit and sequenced on an Illumina MiSeq machine with a 2×150nt sequencing kit, except for the library for strain DEL2, which was sequenced on a Illumina NextSeq 500. Macrogen Inc. (Seoul, South Korea) generated the PacBio data on an RSII machine. The Illumina data was adaptertrimmed using Adapterremoval2 (v. 2.2.2) (64) with the switches --trimns –trimqualities. As described in (34), the genome of *S. griseofuscus* was assembled using PacBio data with the assembly program Flye (v. Flye 2.4.1-geb89c9e) (65) with the switches --genome-size 8m --iterations 5 for five consecutive rounds of polishing using the PacBio data. The assembly was then polished with the Illumina data using the polishing module of Unicycler (v. 0.4.8-beta) (66). After this assembly process, the inverted repeat (IR) chromosome ends were detached from the rest of the chromosome on a separate contig. We manually added the IR to both ends of the chromosome, repeated the illumina polishing with unicycler-polish, and used Minimap2 (v. 2.16-r922) (67) and Bowtie2 (v. 2.3.5) (68) to map pacbio and illumina reads to the manually curated genome assembly to confirm that the IR ends of the chromosome were correctly attached. We used artemis genome viewer (v. 18.0.2) (69) to visualize the mappings. BUSCO (v. 4.0.5) (70) was used for estimating the quality of the genome assembly and Bandage (v. 0.8.1) (71) was used to view and evaluate the assembly graph. The assembled genome was gene annotated using Prodigal (v.2.6.3) (72) and the identified genes were functionally annotated using Prokka (v. 1.14.0) (73) with the PFAM-A (v. 32.0) (74) database and six publically available manually annotated actinobacterial genomes of high quality using the prokka --proteins switch (see Table SI6). These databases were used in addition to the default databases, not instead of them. RNAmmer (75) was used for rRNA gene prediction and Aragorn was used for tRNA prediction (76). Nanopore data from the strain DEL2 was obtained to check for circularization of the genome. The DNA of the DEL2 strain was extracted as described above, and a nanopore library was constructed with the rapid (SQK-RBK004) kit from Oxford Nanopore Technologies (Oxford, United Kingdom) and the barcode RB05. The data was generated on a MinION machine with a 9.4.1 flowcell. The raw data was demultiplexed using Deepbinner (v. 0.2.0) (77) and basecalled using Guppy (v. 3.2.2+9fe0a78, Oxford Nanopore Technologies, Oxford, United Kingdom), before the technical sequence was removed using Porechop (v. 0.2.4; https://github.com/rrwick/Porechop). The Nanopore data was assembled using flye assembler as described above for PacBio data.

### 7. Comparison of BGCs of *S. griseofuscus* with BGCs detected across *Streptomyces sp*

Genome sequences with complete assembly annotation of *Streptomycetaceae* group are collected from the PATRIC database (78) Among 214 complete genome sequences, one was annotated of poor quality owing to its low fine consistency score (PATRIC ID: 2588708.3). One other genome had 22 contigs (PATRIC ID: 1969.5). Remaining 212 genomes of high quality are processed with antiSMASH v5.1 to detect 6380 BGCs in total. The antiSMASH predicted 34 BGCs on chromosome and one BGC on plasmid of *S. griseofuscus*. Some of these BGCs were further split into candidate clusters manually. We used BiG-SCAPE software (79) to generate similarity network of all 6380 BGCs detected in 212 public genomes, 1808 known BGCs from MIBIG database v1.3, 35 BGCs from *S. griseofuscus* genomes and manually selected 14 candidate clusters. The cutoffs of 0.3, 0.5 and 0.7 were used on raw_index similarity metric of BiG-SCAPE analysis. The similarity network was visualized using Cytoscape v3.7. Further some of the candidate clusters of interest (e.g. candidate cluster number 5) were analyzed using the CORASON feature of BiG-SCAPE to generate gene cluster alignments based on similarity.

### 8. Extractions and evaluation of metabolite production

For targeted and untargeted metabolomics analysis, 50 ml liquid cultures with the medium of interest (ISP2 and CDMS) were prepared and cultivated in 250 ml shake flasks for 5 days. After 5 days, the cultures were harvested and centrifuged for 20 min at 10,000 x g. The supernatants were collected in 250 ml glass bottles and extracted using equal volumes (1:1) of ethyl acetate (liquid-liquid extraction). The organic phase was separated from the aqueous phase using a 500 ml separatory funnel, after shortly shaking, and collected in a 250 ml glass bottle. The process was repeated 3 times using fresh solvent. The extracts were combined for each sample and evaporated using a Büchi Rotavapor R-300 in combination with a Büchi Heating Bath B-300 Base, a Büchi Interface I-300, a Büchi Vacuum Pump V-300, and a Julaba Recirculating Cooler FL601. The temperature of the water bath was set to 38 °C, the rotation speed to 140 rpm, and the pressure to 150 mbar, which was gradually decreased to 50 mbar to complete dryness. Next, the analytes were dissolved in 1 ml of 50% v/v methanol and transferred to 2.0 mL Eppendorf tubes. The samples were centrifuged for 10 minutes in 12,000 CFU and the supernatants were transferred in new eppendorf tubes. The samples were again evaporated using an Eppendorf Concentrator Plus Speedvac without heating, dissolved in 100 µl of 50% v/v methanol and centrifuged for 10 min in 12,000 CFU. The supernatants were transferred to autosampler vials with inserts, and analyzed using a LC-MS system. Analysis was performed using a Thermo Dionex Ultimate 3000 UHPLC system with a diode array detector (DAD) interfaced with an Orbitrap Fusion Tribid mass spectrometer (Thermo Scientific, San Jose, USA), using an EASY-IC ESI source. Separation conditions were as follows: Column, Agilent Zorbax Eclipse Plus C18, 100 x 2.1 mm i.d., 1.8 µm particles. The mobile phase used was (A) purified water with 0.1 % formic acid and (B) acetonitrile with 0.1 % formic acid. The flowrate was set to 0.35 ml/min and column temperature was set to 35 °C, while the injection volume was set to 1 µl. The following gradient was used: 0-0.5 min. 5 % B, increasing to 100 % B at 13 min., and holding until 15 min., returning to 5 % B at 15.1 min., and equilibrating for 1.9 min., for a total run time of 17 min. Full-scan mass spectrometric detection was performed in positive and negative ESI mode with the following parameters: source voltage, 3500 V (positive mode) and 2700 V (negative mode); sheath gas flow rate (N2), 50 arbitrary units; auxiliary gas, 10 arbitrary units; sweep gas, 1 arbitrary unit; Ion transfer tube temperature, 325 °C; Vaporizer temperature, 350 °C. MS full-scan analysis was performed using the orbitrap with the following settings: Orbitrap resolution, 120,000; scan range 100-1,000 Da; RF lens, 50 %. Before analysis, the MS was calibrated using ESI Positive Ion Calibration Solution (P/N 88323) and ESI Negative on Calibration Solution (P/N 88324, Thermo Scientific, San Jose, USA). Fragmentation data for compound annotation were obtained using data-dependent MS/MS analysis by selecting the top four most intense ions per cycle. Dynamic exclusion was used to exclude ions for 3 s after two measurements within 4 s. Fragmentation was performed using an assisted fragmentation HCD 15, 30, 45, and 60 % at a resolution of 30,000 with an AGC target of 1×10^5^ and a maximum injection time of 64 ms. Data analysis was performed using Thermo Scientific Compound Discoverer 3.0.0.294. Using the software, data from the LC-MS analysis were aligned, and compound annotation was performed by matching against molecular formulas from StreptomeDB2 and AntiBase, as well as fragmentation spectra from mzCloud. In the case of pentamycin detection, a pure pentamycin standard was used, diluted in 10% v/v methanol to 10^−3^ mg/ml concentration. In regards to the other compounds, no standards were measured, due to their absence in the market, so their detection is putative.

## Data availability

The Genbank accession numbers for *S. griseofuscus* genome are <u>CP051006</u> (chromosome), <u>CP051007</u> (pSGRIFU1), <u>CP051008</u> (pSGRIFU2), and <u>CP051009</u> (pSGRIFU3). All sequencing data can be found in BioProject <u>PRJNA622435</u>.

Following data files including Excel tables and XML format model files are available at link : (https://figshare.com/s/ad8ff9010782c1bc1032).

Dataset 1. Genome comparison data Excel tables with gene annotations from NCBI genbank and KEGG subsystems. Gene presence-absence table for genes across the pangenome of S. griseofuscus, S. coelicolor, S. venezuelae. KEGG subsystem gene and reaction counts per organism.

Dataset 2. Phenotype microarray data analysis with DuctApe Excel tables with raw kinetic growth data across plates PM1 to PM4 with 2 replicates for S. griseofuscus, S. coelicolor, S. venezuelae. Biolog activity index (0 to 9) across 384 nutrients in each strain and comparison of number of active nutrients across the three strains.

Dataset 3. Genotype phenotype mapping using KEGG pathways Excel tables with matrix of biolog activity index across different KEGG nutrients mapped on to the KEGG pathways. Combined matrix with mean activity differences across three strains.

Dataset 4. Genome scale metabolic models The SMBL formatted genome-scale metabolic models of the three *Streptomyces* strains used in the study

Dataset 5: Comparison of genome scale model based growth predictions and BioLog data Excel table with list of reactions considered in the three genome scale models and comparison of in-silico growth prediction on different nutrients against BioLog data.

Dataset 6. BGC comparison across *Streptomyces* using BiGSCAPE Information on BGCs detected in *S. griseofuscus* and information on the public genomes selected along with the BGCs selected for comparison. Similarity network data used for generation of Fig. 3 with information on nodes and edges.

## Legends of Supplementary Information files

Fig. S1. Phenotype of *S. griseofuscus* strains on solid media. 1 - ISP2 media; 2 - MS media; 3 - ISP4 media. A-wild type; B - *S. griseofuscus* IppsD_1 (p054); C - *S. griseofuscus* ItycC_2 (p056); D -*S. griseofuscus* HEP81_06602 (p057); E - *S. griseofuscus* IspkC (p059); F - *S. griseofuscus* DEL1; G - *S. griseofuscus* DEL2; I - *S. griseofuscus* E3I2

Fig. S2. OD600 measurements of *S. griseofuscus*-derived strains, while grown in ISP2 liquid media. A - characterization of DEL1, DEL2, E3I2 strains in comparison to the wild type; B - characterization of slow growing strain p056 in comparison to the wild type; C - characterization of *S. griseofuscus* IppsD_1 (p054), *S. griseofuscus* HEP81_06602 (p057) and *S. griseofuscus* IspkC (p059) strains in comparison to the wild type; D - comparison of all 3 independant cultivations of wild type strains

Fig. S3. Alignment of selected families of BGCs. Here selected regions and candidate clusters of *S. griseofuscus* are aligned against similar BGCs detected across the dataset. Note: Pentamycin BGC was manually added in the above analysis as default BiGSCAPE uses an older MIBIG version lacking pentamycin BGC.

Fig. S4. Results of metabolite production analysis of cultivations and extractions of wild type, DEL1 and DEL2 strains, grown in liquid ISP2 media. A) Extracted ion chromatograms of pentamycin (C35H58O12) production of strains WT, DEL1 and DEL2 cultivated in ISP2 media in comparison to the extraction of pure ISP2 media and pentamycin standard. All cultivations were done in triplicates; B) the extracted EICs for kujimycin A (C40H70O15) are plotted for comparative metabolomics between blank ISP2 medium, three replicates of *S. griseofuscus* WT in ISP2 and three replicates of *S. griseofuscus* DEL1 in ISP2. The peak at 8.70 min is a putative match for the molecule and it is clearly absent in case of DEL1 strain

Fig. S5. A and B - three independent colonies of *S. griseofuscus* pKG1139 (I), *S. griseofuscus* pSETGUS (II) and wild type (III), grown on ISP2 solid media after the 24 hours incubation at 40°C to verify possibility of plasmid curing, A-colonies before the addition of X-Gluc; B - colonies after the addition of X-Gluc

Fig. S6. Strains *S. griseofuscus* wild type, DEL1 and DEL2, that contain integrative plasmid construct, carrying complete actinorhodin gene cluster, cloned from *S. coelicolor* M145. All strains were grown on MS media, supplemented with apramycin for 6 days before the photo was taken

Table S1. OD600 measurements of *S. griseofuscus*-derived strains, while grown in ISP2 liquid media. The growth curve was built in liquid ISP2 culture based on three independently performed cultivations, each of which consisted of three cultivation flasks and each data point was measured three times. Please, review this table in connection to Fig. S3. The µ_max_ for the wild type was calculated to be 0.43 ± 0.08 h^-1^ on average for three cultivations with td of 1.68 ± 0.29 h. Sporulation of the DEL1 and DEL2 strains was estimated for their growth on MS media, where the wild type sporulation was accounted for at 3,43±3,05×10^8^ CFU/mL. Wild type sporulation was observed only on MS and ISP4 media, but not ISP2, with an average amount of spores on ISP4 - 2,60±4,58×10^8^ CFU/mL, therefore MS media was chosen as the most suitable for growth experiments.

Table S2. Compounds detected in ethyl acetate extractions of supernatants of *S. griseofuscus* wild type cultivations in different liquid media. RT - retention time, MM - minimal medium, M65 - medium 65. Detection of pentamycin was confirmed via comparison with a pure pentamycin standard

Table S3. BLAST hits of genes *tapR1/tpgR1* from *S. rochei* 7434AN4 genome to genes in *S. griseofuscus* genome

Table S4. Summary of all primers, genetic constructs and strains used in this study

## Acknowledgements

The authors would like to thank Alexandra Hoffmeyer for her invaluable help with sequencing, Andreas Klitgaard and Yulia Radko for the help with analyzing LCMS data, Kai Blin for the help with bioinformatic analysis, Yaojun Tong for his help with CRISPR-Cas9 related experiments, Xinglin Jiang for the help with cloning experiments.

The work of the authors is funded by grants of the Novo Nordisk Foundation, Denmark [NNF10CC1016517, NNF16OC0021746].

## References

1. Hutchings MI, Truman AW, Wilkinson B. 2019. Antibiotics: past, present and future. Curr Opin Microbiol 51:72–80.

2. Genilloud O. 2017. Actinomycetes: still a source of novel antibiotics. Nat Prod Rep 34:1203–1232.

3. Blin K, Shaw S, Steinke K, Villebro R, Ziemert N, Lee SY, Medema MH, Weber T. 2019. antiSMASH 5.0: updates to the secondary metabolite genome mining pipeline. Nucleic Acids Res 47:W81–W87.

4. Belknap KC, Park CJ, Barth BM, Andam CP. 2020. Genome mining of biosynthetic and chemotherapeutic gene clusters in Streptomyces bacteria. 1. Sci Rep 10:1–9.

5. Hoskisson PA, Seipke RF. 2020. Cryptic or Silent? The Known Unknowns, Unknown Knowns, and Unknown Unknowns of Secondary Metabolism. mBio 11.

6. Tomm HA, Ucciferri L, Ross AC. 2019. Advances in microbial culturing conditions to activate silent biosynthetic gene clusters for novel metabolite production. J Ind Microbiol Biotechnol 46:1381–1400.

7. Chen C, Zhao X, Chen L, Jin Y, Zhao ZK, Suh J-W. 2015. Effect of overexpression of endogenous and exogenous Streptomyces antibiotic regulatory proteins on tacrolimus (FK506) production in Streptomyces sp. KCCM11116P. RSC Adv 5:15756–15762.

8. Wei Q, Aung A, Liu B, Ma J, Shi L, Zhang K, Ge B. 2020. Overexpression of wysR gene enhances wuyiencin production in ΔwysR3 mutant strain of Streptomyces albulus var. wuyiensis strain CK-15. J Appl Microbiol 129:565–574.

9. Wang B, Guo F, Dong S-H, Zhao H. 2019. Activation of silent biosynthetic gene clusters using transcription factor decoys. 2. Nat Chem Biol 15:111–114.

10. Horbal L, Marques F, Nadmid S, Mendes MV, Luzhetskyy A. 2018. Secondary metabolites overproduction through transcriptional gene cluster refactoring. Metab Eng 49:299–315.

11. Palazzotto E, Tong Y, Lee SY, Weber T. 2019. Synthetic biology and metabolic engineering of actinomycetes for natural product discovery. Biotechnol Adv 37:107366.

12. Culp EJ, Yim G, Waglechner N, Wang W, Pawlowski AC, Wright GD. 2019. Hidden antibiotics in actinomycetes can be identified by inactivation of gene clusters for common antibiotics. Nat Biotechnol 37:1149–1154.

13. Hug JJ, Krug D, Müller R. 2020. Bacteria as genetically programmable producers of bioactive natural products. Nat Rev Chem 1–22.

14. Myronovskyi M, Luzhetskyy A. 2019. Heterologous production of small molecules in the optimized Streptomyces hosts. Nat Prod Rep 36:1281–1294.

15. Ahmed Y, Rebets Y, Estévez MR, Zapp J, Myronovskyi M, Luzhetskyy A. 2020. Engineering of Streptomyces lividans for heterologous expression of secondary metabolite gene clusters. Microb Cell Factories 19:5.

16. Bu Q-T, Yu P, Wang J, Li Z-Y, Chen X-A, Mao X-M, Li Y-Q. 2019. Rational construction of genome-reduced and high-efficient industrial Streptomyces chassis based on multiple comparative genomic approaches. Microb Cell Factories 18:16.

17. Ikeda H, Kazuo S, Omura S. 2014. Genome mining of the Streptomyces avermitilis genome and development of genome-minimized hosts for heterologous expression of biosynthetic gene clusters. J Ind Microbiol Biotechnol 41:233–250.

18. Komatsu M, Komatsu K, Koiwai H, Yamada Y, Kozone I, Izumikawa M, Hashimoto J, Takagi M, Omura S, Shin-ya K, Cane DE, Ikeda H. 2013. Engineered Streptomyces avermitilis host for heterologous expression of biosynthetic gene cluster for secondary metabolites. ACS Synth Biol 2:384–396.

19. Kim EJ, Yang I, Yoon YJ. 2015. Developing Streptomyces venezuelae as a cell factory for the production of small molecules used in drug discovery. Arch Pharm Res 38:1606–1616.

20. Yin S, Li Z, Wang X, Wang H, Jia X, Ai G, Bai Z, Shi M, Yuan F, Liu T, Wang W, Yang K. 2016. Heterologous expression of oxytetracycline biosynthetic gene cluster in Streptomyces venezuelae WVR2006 to improve production level and to alter fermentation process. Appl Microbiol Biotechnol 100:10563–10572.

21. Myronovskyi M, Rosenkränzer B, Nadmid S, Pujic P, Normand P, Luzhetskyy A. 2018. Generation of a cluster-free Streptomyces albus chassis strains for improved heterologous expression of secondary metabolite clusters. Metab Eng 49:316–324.

22. Fazal A, Thankachan D, Harris E, Seipke RF. 2020. A chromatogram-simplified Streptomyces albus host for heterologous production of natural products. Antonie Van Leeuwenhoek 113:511–520.

23. Myronovskyi M, Rosenkränzer B, Stierhof M, Petzke L, Seiser T, Luzhetskyy A. 2020. Identification and Heterologous Expression of the Albucidin Gene Cluster from the Marine Strain Streptomyces Albus Subsp. Chlorinus NRRL B-24108. Microorganisms 8.

24. Rodríguez Estévez M, Gummerlich N, Myronovskyi M, Zapp J, Luzhetskyy A. 2019. Benzanthric Acid, a Novel Metabolite From Streptomyces albus Del14 Expressing the Nybomycin Gene Cluster. Front Chem 7:896.

25. Sakamoto JM, Kondo SI, Yumoto H, Arishima M. 1962. Bundlins A and B, two antibiotics produced by Strepto. J Antibiot (Tokyo) 15:98–102.

26. Larson JL, Hershberger CL. 1984. Shuttle vectors for cloning recombinant DNA in Escherichia coli and Streptomyces griseofuscus C581. J Bacteriol 157:314–317.

27. Larson JL, Hershberger CL. 1986. The minimal replicon of a streptomycete plasmid produces an ultrahigh level of plasmid DNA. Plasmid 15:199–209.

28. McHenney MA, Baltz RH. 1988. Transduction of plasmid DNA in Streptomyces spp. And related genera by bacteriophage FP43. J Bacteriol 170:2276–2282.

29. Li S, Tang L, Chen X, Liao L, Li F, Mao Z. 2011. Isolation and characterization of a novel ε-poly-l-lysine producing strain: Streptomyces griseofuscus. J Ind Microbiol Biotechnol 38:557–563.

30. Li S, Chen X, Dong C, Zhao F, Tang L, Mao Z. 2013. Combining Genome Shuffling and Interspecific Hybridization Among Streptomyces Improved ε-Poly-l-Lysine Production. Appl Biochem Biotechnol 169:338–350.

31. Li S, Ji J, Hu S, Chen G. 2020. Enhancement of ε-poly-L-lysine production in Streptomyces griseofuscus by addition of exogenous astaxanthin. Bioprocess Biosyst Eng 43:1813–1821.

32. Lacalle RA, Tercero JA, Jiménez A. 1992. Cloning of the complete biosynthetic gene cluster for an aminonucleoside antibiotic, puromycin, and its regulated expression in heterologous hosts. EMBO J 11:785–792.

33. Wang B, Guo F, Huang C, Zhao H. 2020. Unraveling the iterative type I polyketide synthases hidden in Streptomyces. Proc Natl Acad Sci U S A 117:8449–8454.

34. Gren T, Jørgensen T, Whitford C, Weber T. 2020. High-quality sequencing, assembly and annotation of the Streptomyces griseofuscus DSM 40191 genome. Microbiol Resour Announc Press.

35. Moriya Y, Itoh M, Okuda S, Yoshizawa AC, Kanehisa M. 2007. KAAS: an automatic genome annotation and pathway reconstruction server. Nucleic Acids Res 35:W182–W185.

36. Koepff J, Keller M, Tsolis KC, Busche T, Rückert C, Hamed MB, Anné J, Kalinowski J, Wiechert W, Economou A, Oldiges M. 2017. Fast and reliable strain characterization of Streptomyces lividans through micro-scale cultivation. Biotechnol Bioeng 114:2011–2022.

37. Galardini M, Mengoni A, Biondi EG, Semeraro R, Florio A, Bazzicalupo M, Benedetti A, Mocali S. 2014. DuctApe: A suite for the analysis and correlation of genomic and OmniLogTM Phenotype Microarray data. Genomics 103:1–10.

38. Medema MH, Kottmann R, Yilmaz P, Cummings M, Biggins JB, Blin K, de Bruijn I, Chooi YH, Claesen J, Coates RC, Cruz-Morales P, Duddela S, Düsterhus S, Edwards DJ, Fewer DP, Garg N, Geiger C, Gomez-Escribano JP, Greule A, Hadjithomas M, Haines AS, Helfrich EJN, Hillwig ML, Ishida K, Jones AC, Jones CS, Jungmann K, Kegler C, Kim HU, Kötter P, Krug D, Masschelein J, Melnik AV, Mantovani SM, Monroe EA, Moore M, Moss N, Nützmann H-W, Pan G, Pati A, Petras D, Reen FJ, Rosconi F, Rui Z, Tian Z, Tobias NJ, Tsunematsu Y, Wiemann P, Wyckoff E, Yan X, Yim G, Yu F, Xie Y, Aigle B, Apel AK, Balibar CJ, Balskus EP, Barona-Gómez F, Bechthold A, Bode HB, Borriss R, Brady SF, Brakhage AA, Caffrey P, Cheng Y-Q, Clardy J, Cox RJ, De Mot R, Donadio S, Donia MS, van der Donk WA, Dorrestein PC, Doyle S, Driessen AJM, Ehling-Schulz M, Entian K-D, Fischbach MA, Gerwick L, Gerwick WH, Gross H, Gust B, Hertweck C, Höfte M, Jensen SE, Ju J, Katz L, Kaysser L, Klassen JL, Keller NP, Kormanec J, Kuipers OP, Kuzuyama T, Kyrpides NC, Kwon H-J, Lautru S, Lavigne R, Lee CY, Linquan B, Liu X, Liu W, Luzhetskyy A, Mahmud T, Mast Y, Méndez C, Metsä-Ketelä M, Micklefield J, Mitchell DA, Moore BS, Moreira LM, Müller R, Neilan BA, Nett M, Nielsen J, O’Gara F, Oikawa H, Osbourn A, Osburne MS, Ostash B, Payne SM, Pernodet J-L, Petricek M, Piel J, Ploux O, Raaijmakers JM, Salas JA, Schmitt EK, Scott B, Seipke RF, Shen B, Sherman DH, Sivonen K, Smanski MJ, Sosio M, Stegmann E, Süssmuth RD, Tahlan K, Thomas CM, Tang Y, Truman AW, Viaud M, Walton JD, Walsh CT, Weber T, van Wezel GP, Wilkinson B, Willey JM, Wohlleben W, Wright GD, Ziemert N, Zhang C, Zotchev SB, Breitling R, Takano E, Glöckner FO. 2015. Minimum Information about a Biosynthetic Gene cluster. 9. Nat Chem Biol 11:625–631.

39. Jain C, Rodriguez-R LM, Phillippy AM, Konstantinidis KT, Aluru S. 2018. High throughput ANI analysis of 90K prokaryotic genomes reveals clear species boundaries. 1. Nat Commun 9:5114.

40. Cao Z, Yoshida R, Kinashi H, Arakawa K. 2015. Blockage of the early step of lankacidin biosynthesis caused a large production of pentamycin, citreodiol and epi -citreodiol in Streptomyces rochei. 5. J Antibiot (Tokyo) 68:328–333.

41. Nindita Y, Cao Z, Fauzi AA, Teshima A, Misaki Y, Muslimin R, Yang Y, Shiwa Y, Yoshikawa H, Tagami M, Lezhava A, Ishikawa J, Kuroda M, Sekizuka T, Inada K, Kinashi H, Arakawa K. 2019. The genome sequence of Streptomyces rochei 7434AN4, which carries a linear chromosome and three characteristic linear plasmids. Sci Rep 9:10973.

42. Arakawa K. 2014. Genetic and biochemical analysis of the antibiotic biosynthetic gene clusters on the Streptomyces linear plasmid. Biosci Biotechnol Biochem 78:183–189.

43. Nagaoka K, Matsumoto M, Oono J, Yokoi K, Ishizeki S, Nakashima T. 1986. Azinomycins A and B, new antitumor antibiotics. I. Producing organism, fermentation, isolation, and characterization. J Antibiot (Tokyo) 39:1527–1532.

44. Zhang J, Marcin C, Shifflet MA, Salmon P, Brix T, Greasham R, Buckland B, Chartrain M. 1996. Development of a defined medium fermentation process for physostigmine production by Streptomyces griseofuscus. Appl Microbiol Biotechnol 44:568–575.

45. Kieser T, Bibb MJ, Buttner MJ, Chater KF, Hopwood DA. 2000. Practical streptomyces genetics. John Innes Foundation Norwich.

46. Gym Streptomyces medium.

47. Fröjd MJ, Flärdh K. 2019. Extrusion of extracellular membrane vesicles from hyphal tips of Streptomyces venezuelae coupled to cell-wall stress. Microbiology 165:1295–1305.

48. Hoefler BC, Stubbendieck RM, Josyula NK, Moisan SM, Schulze EM, Straight PD. 2017. A Link between Linearmycin Biosynthesis and Extracellular Vesicle Genesis Connects Specialized Metabolism and Bacterial Membrane Physiology. Cell Chem Biol 24:1238-1249.e7.

49. Schrempf H, Merling P. 2015. Extracellular Streptomyces lividans vesicles: composition, biogenesis and antimicrobial activity. Microb Biotechnol 8:644–658.

50. Tercero JA, Lacalle RA, Jiménez A. 1992. Cosmid pJAR4, a novel Streptomyces-Escherichia coli shuttle vector for the cloning of Streptomyces operons. FEMS Microbiol Lett 75:203–206.

51. Myronovskyi M, Welle E, Fedorenko V, Luzhetskyy A. 2011. Beta-glucuronidase as a sensitive and versatile reporter in actinomycetes. Appl Environ Microbiol 77:5370–5383.

52. Gren T, Ortseifen V, Wibberg D, Schneiker-Bekel S, Bednarz H, Niehaus K, Zemke T, Persicke M, Pühler A, Kalinowski J. 2016. Genetic engineering in Actinoplanes sp. SE50/110 - development of an intergeneric conjugation system for the introduction of actinophage-based integrative vectors. J Biotechnol 232:79–88.

53. Baltz RH. 2012. Streptomyces temperate bacteriophage integration systems for stable genetic engineering of actinomycetes (and other organisms). J Ind Microbiol Biotechnol 39:661–672.

54. Tong Y, Weber T, Lee SY. 2019. CRISPR/Cas-based genome engineering in natural product discovery. Nat Prod Rep 36:1262–1280.

55. Tong Y, Charusanti P, Zhang L, Weber T, Lee SY. 2015. CRISPR-Cas9 Based Engineering of Actinomycetal Genomes. ACS Synth Biol 4:1020–1029.

56. Nindita Y, Cao Z, Yang Y, Arakawa K, Shiwa Y, Yoshikawa H, Tagami M, Lezhava A, Kinashi H. 2015. The tap-tpg gene pair on the linear plasmid functions to maintain a linear topology of the chromosome in Streptomyces rochei. Mol Microbiol 95:846–858.

57. Tong Y, Whitford CM, Robertsen HL, Blin K, Jørgensen TS, Klitgaard AK, Gren T, Jiang X, Weber T, Lee SY. 2019. Highly efficient DSB-free base editing for streptomycetes with CRISPR-BEST. Proc Natl Acad Sci U S A 116:20366–20375.

58. Hoff G, Bertrand C, Piotrowski E, Thibessard A, Leblond P. 2018. Genome plasticity is governed by double strand break DNA repair in Streptomyces. Sci Rep 8:5272.

59. Tong Y, Whitford CM, Blin K, Jørgensen TS, Weber T, Lee SY. 2020. CRISPR-Cas9, CRISPRi and CRISPR-BEST-mediated genetic manipulation in streptomycetes. Nat Protoc 15:2470–2502.

60. Ye S, Enghiad B, Zhao H, Takano E. 2020. Fine-tuning the regulation of Cas9 expression levels for efficient CRISPR-Cas9 mediated recombination in Streptomyces. J Ind Microbiol Biotechnol 47:413–423.

61. 2013. Copyright, p. ii. edIn Doran, PM (ed.), Bioprocess Engineering Principles (Second Edition). Academic Press, London.

62. Blin K, Pedersen LE, Weber T, Lee SY. 2016. CRISPy-web: An online resource to design sgRNAs for CRISPR applications. Synth Syst Biotechnol 1:118–121.

63. Blin K, Shaw S, Tong Y, Weber T. 2020. Designing sgRNAs for CRISPR-BEST base editing applications with CRISPy-web 2.0. Synth Syst Biotechnol 5:99–102.

64. Schubert M, Lindgreen S, Orlando L. 2016. AdapterRemoval v2: rapid adapter trimming, identification, and read merging. BMC Res Notes 9:88.

65. Kolmogorov M, Yuan J, Lin Y, Pevzner PA. 2019. Assembly of long, error-prone reads using repeat graphs. Nat Biotechnol 37:540–546.

66. Wick RR, Judd LM, Gorrie CL, Holt KE. 2017. Unicycler: Resolving bacterial genome assemblies from short and long sequencing reads. PLoS Comput Biol 13:e1005595.

67. Li H. 2018. Minimap2: pairwise alignment for nucleotide sequences. Bioinformatics 34:3094–3100.

68. Langmead B, Salzberg SL. 2012. Fast gapped-read alignment with Bowtie 2. 4. Nat Methods 9:357–359.

69. Carver T, Harris SR, Berriman M, Parkhill J, McQuillan JA. 2012. Artemis: an integrated platform for visualization and analysis of high-throughput sequence-based experimental data. Bioinformatics 28:464–469.

70. Seppey M, Manni M, Zdobnov EM. 2019. BUSCO: Assessing Genome Assembly and Annotation Completeness, p. 227–245. In Kollmar, M (ed.), Gene Prediction: Methods and Protocols. Springer, New York, NY.

71. Wick RR, Schultz MB, Zobel J, Holt KE. 2015. Bandage: interactive visualization of de novo genome assemblies. Bioinforma Oxf Engl 31:3350–3352.

72. Hyatt D, Chen G-L, Locascio PF, Land ML, Larimer FW, Hauser LJ. 2010. Prodigal: prokaryotic gene recognition and translation initiation site identification. BMC Bioinformatics 11:119.

73. Seemann T. 2014. Prokka: rapid prokaryotic genome annotation. Bioinforma Oxf Engl 30:2068–2069.

74. El-Gebali S, Mistry J, Bateman A, Eddy SR, Luciani A, Potter SC, Qureshi M, Richardson LJ, Salazar GA, Smart A, Sonnhammer ELL, Hirsh L, Paladin L, Piovesan D, Tosatto SCE, Finn RD. 2019. The Pfam protein families database in 2019. Nucleic Acids Res 47:D427–D432.

75. Lagesen K, Hallin P, Rødland EA, Staerfeldt H-H, Rognes T, Ussery DW. 2007. RNAmmer: consistent and rapid annotation of ribosomal RNA genes. Nucleic Acids Res 35:3100–3108.

76. Laslett D, Canback B. 2004. ARAGORN, a program to detect tRNA genes and tmRNA genes in nucleotide sequences. Nucleic Acids Res 32:11–16.

77. Wick RR, Judd LM, Holt KE. 2018. Deepbinner: Demultiplexing barcoded Oxford Nanopore reads with deep convolutional neural networks. PLOS Comput Biol 14:e1006583.

78. Davis JJ, Wattam AR, Aziz RK, Brettin T, Butler R, Butler RM, Chlenski P, Conrad N, Dickerman A, Dietrich EM, Gabbard JL, Gerdes S, Guard A, Kenyon RW, Machi D, Mao C, Murphy-Olson D, Nguyen M, Nordberg EK, Olsen GJ, Olson RD, Overbeek JC, Overbeek R, Parrello B, Pusch GD, Shukla M, Thomas C, VanOeffelen M, Vonstein V, Warren AS, Xia F, Xie D, Yoo H, Stevens R. 2020. The PATRIC Bioinformatics Resource Center: expanding data and analysis capabilities. Nucleic Acids Res 48:D606–D612.

79. Navarro-Muñoz JC, Selem-Mojica N, Mullowney MW, Kautsar SA, Tryon JH, Parkinson EI, De Los Santos ELC, Yeong M, Cruz-Morales P, Abubucker S, Roeters A, Lokhorst W, Fernandez-Guerra A, Cappelini LTD, Goering AW, Thomson RJ, Metcalf WW, Kelleher NL, Barona-Gomez F, Medema MH. 2020. A computational framework to explore large-scale biosynthetic diversity. Nat Chem Biol 16:60–68.

